# Spatial association between distributed β-amyloid and tau varies with cognition

**DOI:** 10.1101/2023.09.27.559737

**Authors:** Felix Carbonell, Carolann McNicoll, Alex P. Zijdenbos, Barry J. Bedell, Alzheimer’s Disease Neuroimaging Initiative

## Abstract

Several PET studies have explored the relationship between β-amyloid load and tau uptake at the early stages of Alzheimer’s disease (AD) progression. Most of these studies have focused on the linear relationship between β-amyloid and tau at the local level and their synergistic effect on different AD biomarkers. We hypothesize that patterns of spatial association between β-amyloid and tau might be uncovered using alternative association metrics that account for linear as well as more complex, possible nonlinear dependencies. In the present study, we propose a new Canonical Distance Correlation Analysis (CDCA) to generate distinctive spatial patterns of the cross-correlation structure between tau, as measured by [18F]flortaucipir PET, and β-amyloid, as measured by [18F]florbetapir PET, from the Alzheimer’s Disease Neuroimaging Initiative (ADNI) study. We found that the CDCA-based β-amyloid scores were not only maximally distance-correlated to tau in cognitively normal (CN) controls and mild cognitive impairment (MCI), but also differentiated between low and high levels of β-amyloid uptake. The most distinctive spatial association pattern was characterized by a spread of β-amyloid covering large areas of the cortex and localized tau in the entorhinal cortex. More importantly, this spatial dependency varies according to cognition, which cannot be explained by the uptake differences in β-amyloid or tau between CN and MCI subjects. Hence, the CDCA-based scores might be more accurate than the amyloid or tau SUVR for the enrollment in clinical trials of those individuals on the path of cognitive deterioration.

## Introduction

Widespread accumulation of β-amyloid plaques and neurofibrillary tangles comprised of tau aggregates are considered prominent neuropathologic features of Mild Cognitive Impairment (MCI) and Alzheimer’s disease (AD)[1–4]. The early consensus about the AD pathogenesis relied on the so-called “amyloid cascade hypothesis” [5–7] that focused the interest on the β-amyloid deposition process as the key triggering event that induces a cascade of time-ordered events including tau pathology, glucose hypometabolism, brain atrophy, cognitive impairment, and dementia. As such, investigators have extensively tested the amyloid cascade hypothesis to gain a better understanding of the relationship between β-amyloid deposition and AD-associated pathophysiological changes, such as tau accumulation and, ultimately, neuronal loss and cognitive deterioration [7–14]. Those early studies primarily focused on β-amyloid deposition as the triggering mechanism of AD, while associating the tau pathology with later events leading to neuronal loss. However, more recent studies support the alternative hypothesis that the accumulation of β-amyloid plaques and neurofibrillary tangles composed of tau aggregates more likely act as coordinated and parallel events in the course of the disease progression [15–17]. The amount of evidence questioning the hypothesis about the triggering role of β-amyloid together with the accumulation of negative results from several anti-Aβ clinical trials [18–20] has led to the need to reformulate the initial amyloid cascade hypothesis not only from the mechanistic point of view, but also from the therapeutic perspective [18,21–23]. Hence, renewed efforts have been put into gaining a better understanding of the synergetic association between β-amyloid and tau accumulation across the various stages of the disease progression and their impact on other AD biomarkers [16,24–30].

Some studies [24,27,30,31] have focused on characterizing the communalities and differences in the spatial topography of the β-amyloid and tau accumulation in aging populations with different cognitive statuses. These studies attempted to perform an *in vivo* characterization of the proposed Braak stages [3] for the spatiotemporal hierarchical organization of networks of tau accumulation across the AD progression and their relationship with the spread of β-amyloid [31]. Thus, hierarchical clustering techniques in cognitively normal older subjects [24] revealed high tau intensity areas clustered predominantly in the medial temporal lobe, basolateral temporal cortex, and orbitofrontal cortex, while areas showing high β-amyloid burden were mainly located in the lateral temporoparietal and frontal regions. Similarly, Brier et al. [30] used SVD techniques to demonstrate that tau and β-amyloid showed distinctive spatial topographic patterns of accumulation across the cerebral cortex, where tau deposition was demonstrated to be more strongly associated with cognitive performance than β-amyloid.

Nevertheless, the spatial characterization of *in vivo* staging of β-amyloid and tau aggregation has resulted in an insufficient understanding of the synergistic role of these two mechanisms on the rest of AD biomarkers, such as metabolic decline, atrophy, neurodegeneration, and cognitive decline. With this aim, [16,28,32] tested the interactive effects of β-amyloid and CSF-phosphorylated tau (p-tau) on the metabolic decline and cognitive deterioration in preclinical AD. It was demonstrated that the simultaneous interaction of β-amyloid and p-tau, rather than the sum of their independent effects, was associated with a metabolic decline in basal and mesial temporal, orbitofrontal, and anterior and posterior cingulate cortices [16,32], and with cognitive decline and progression to AD [28]. Similarly, [26] investigated the impact of the interaction of β-amyloid and tau pathologies on cognitive decline in normal aging and preclinical AD. Both ROI and voxelwise analysis in [26] revealed that a significant interaction between β-amyloid and tau was strongly associated with the fast memory deterioration observed in participants showing elevated levels of both pathologies.

Despite the accumulating evidence that highlights the coordinated role of β-amyloid and tau in subsequent neurodegeneration, atrophy, and cognitive decline processes, a more detailed spatial characterization of the proper interaction between these two pathologies is still necessary. Initial studies, like [30] used a linear canonical correlation analysis (CCA) to directly compare the relationship between the spatial PET tau and β-amyloid topographies at the ROI level. The CCA in [30] revealed that the spatial loading of the tau and β-amyloid topographies were modestly correlated, even though the first canonical pair (i.e., projected data) showed a strong correlation. In particular, the PET tau topography was primarily localized in the medial temporal lobe and parietal cortex, and it was highly correlated to the β-amyloid topography with strong spatial loads in the frontal and parietal areas. Indeed, the results of [30] highlighted the importance of studying the distributed-to-distributed view of the spatial relationship between tau and β-amyloid as opposed to the more classical global-to-distributed [33,34], local-to-local, and local-to-distributed views [35–37].

The local-to-local analysis in [37] computed partial correlations between PET images of tau and β-amyloid at corresponding voxels while controlling for age and local (i.e., voxel) atrophy measurements in cognitively normal elderly participants. It was found that tau and β-amyloid were significantly locally correlated in areas of the lateral temporal, frontal, and parietal lobes, precuneus, and posterior regions, with the strongest correlations corresponding to the lateral inferior temporal lobe [37]. Additionally, the local-to-distributed cross-correlation analysis of [37] summarizes (i.e., weighted degree of connectivity) to what extent each seed voxel in one PET modality is significantly correlated to each of the voxels in the other PET modality. For instance, it was found that β-amyloid deposits in several areas, including the lateral-ventral frontal, inferior parietal, and temporal cortices showed several significant correlations with distributed tau patterns [37]. Similarly, bilateral inferior-temporal and entorhinal cortices showed a high number of significant tau-distributed-amyloid correlations [37]. Also, in cognitively normal older subjects, [36] showed local-to-local linear correlations between tau and β-amyloid largely located in the temporal lobes. Additional local-to-distributed significant correlations were identified between β-amyloid in multiple regions of the association cortex and tau temporal cortical ROIs [36]. A multi-modal correlation analysis in [35] also examined the local-to-local and the local-to-distributed views of the β-amyloid-tau relationship in a small sample of cognitively normal and mild AD older participants, where it was found that β-amyloid and tau were positively locally correlated within the parietal and the medial/inferior occipital regions. A potential limitation of these previous studies about the spatial relationship between β-amyloid and tau is that they have been primarily focused on cognitively normal individuals. Thus, a more detailed view of the β-amyloid-tau associations across different stages of the disease remains poorly understood. Additionally, it seems that the local-to-local and local-to-distributed views of the relationship β-amyloid-tau have dominated the field. More importantly, all those studies focused only on linear relationships, while more complex potential nonlinear interactions remain uncovered. To overcome these limitations, we focus on revealing distributed-to-distributed and local-to-distributed nonlinear views of the β-amyloid-tau relationship in a sample of CN and MCI older subjects. With this aim, we employ the concepts of Distance-Correlations [38] and distance-induced Kernel Canonical Correlations Analysis (KCCA) [39]. Essentially, Distance-Correlations is an associative metric between multi-dimensional vectors that generalizes the classical Pearson’s correlations in the sense that it can detect not only linear associations, but also general dependencies [38,40]. Additionally, the distance-induced kernel [41] allows one to perform a KCCA to detect general patterns of the cross-dependency between tau and β-amyloid without the need to specify an *a priori* non-linear functional relationship.

In the current study, we hypothesize that distance-correlations as a general measure of dependency can uncover significant patterns of distributed-to-distributed co-variation in the relationship between β-amyloid and tau across the AD progression. We also hypothesize that the distributed-to-distributed and local-to-distributed patterns of the β-amyloid-tau relationship vary according to the different degrees of cognitive impairment.

## Materials and Methods

### Subjects and Image Acquisition

Data used in the preparation of this article were obtained from the ADNI database (http://adni.loni.usc.edu). The ADNI was launched in 2003 by the National Institute on Aging (NIA), the National Institute of Biomedical Imaging and Bioengineering (NIBIB), the Food and Drug Administration (FDA), private pharmaceutical companies, and non-profit organizations, as a $60 million, 5-year public private partnership, which has since been extended. ADNI is the result of the efforts of many co-investigators from a broad range of academic institutions and private corporations, and subjects have been recruited from over 55 sites across the U.S. and Canada. To date, the ADNI, AND-GO, ADNI-2 and ADNI-3 protocols have recruited over 1,500 adults, ages 55 to 90, to participate in the research, consisting of cognitively normal (CN) older individuals, people with early or late Mild Cognitive Impairment (MCI), and people with early AD. For up-to-date information, see www.adni-info.org.

The subjects of this cross-sectional study consisted of 157 participants from the ADNI study who had available [18F]florbetapir PET, [18F]flortaucipir PET, and 3D T1-weighted anatomical MRI within a time frame of 3 months. CN subjects had Mini-Mental State Exam (MMSE) scores between 24 and 30 inclusively, a Clinical Dementia Rating (CDR) of 0, and did not have depression, MCI, or dementia. Early MCI (EMCI) subjects had MMSE scores between 24 and 30 inclusively, a CDR of 0.5, a reported subjective memory concern, an absence of dementia, an objective memory loss measured by education-adjusted scores on delayed recall of one paragraph from the Wechsler Memory Scale Logical Memory (WMSLM) II, essentially preserved activities of daily living, and no impairment in other cognitive domains. MCI subjects had the same inclusion criteria, except for objective memory loss measured by education adjusted scores on delayed recall of one paragraph from WMSLM II.

A detailed description of the ADNI MRI and PET image acquisition protocols can be found at http://adni.loni.usc.edu/methods. ADNI studies are conducted in accordance with the Good Clinical Practice guidelines, the Declaration of Helsinki, and U.S. 21 CFR Part 50 (Protection of Human Subjects) and Part 56 (Institutional Review Boards), where informed written consent was obtained from all participants at each site.

The [18F]florbetapir PET scans were also classified into high (Aβ_H_) and low (Aβ_L_) amyloid subjects according to their SUVR values in an optimal target ROI (Stat-ROI) including areas of the posterior cingulate cortex, precuneus, and medial frontal cortex, with an associated cutoff of SUVR=1.24. That optimal target ROI and cutoff value for the segregation of subjects into low and high β-amyloid were selected by our data-driven approach that has been previously reported [42].

### Image Processing

MR and PET images were processed using the PIANO™ software package (Biospective Inc., Montreal, Canada). T1-weighted MRI volumes underwent image non-uniformity correction using the N3 algorithm [43], brain masking, linear spatial normalization utilizing a 9-parameter affine transformation, and nonlinear spatial normalization to map individual images from native coordinate space to Montreal Neurological Institute (MNI) reference space using a customized, anatomical MRI template derived from ADNI subjects. The resulting image volumes were segmented into gray matter (GM), white matter (WM), and cerebrospinal fluid (CSF) using an artificial neural network classifier [44] and partial volume estimation [45].

The [18F]florbetapir and [18F]flortaucipir PET images underwent several pre-processing steps, including frame-to-frame linear motion correction, smoothing with scanner-specific blurring kernels to achieve 8mm FWHM [46], and averaging of dynamic frames into a static image. The resulting smoothed PET volumes were linearly registered to the subject’s T1-weighted MRI and, subsequently, spatially normalized to reference space using the linear and nonlinear transformations derived from the anatomical MRI registration. Standardized uptake value ratio (SUVR) maps of the PET images were generated from both [18F]florbetapir and [18F]flortaucipir PET using the gray matter(GM)-masked full Cerebellum as a reference region. To minimize PET off-target binding in non-GM areas, we have restricted our analysis to those voxels in the brain cortex that have been previously masked by a GM mask in the nonlinear template space. Hence, all our voxelwise PET maps as well as statistical maps were projected onto the cortical surface for visualization purposes only.

### Canonical Distance Correlation Analysis

The basic underlying idea of the Canonical Distance Correlation Analysis (CDCA) is to use the relatively new concept of Distance Correlation/Covariance [38] to perform a canonical distance-correlation analysis between two sets of PET images (e.g., Amyloid and Tau PET SUVR images).

Traditionally, Singular Value Decomposition (SVD) techniques and Canonical Correlation Analysis (CCA) have been used to detect networks of linear cross-correlation among different brain imaging modalities [47–51] via Pearson’s correlation as the underlying association measure. Despite being widely used, such linear techniques are not able to detect nonlinear and other complex dependencies, particularly in high-dimensional scenarios, like brain images. To overcome this limitation, nonlinear variants of CCA based on kernels have been proposed [39,52–54]. Kernel CCA (KCCA) can detect instances of nonlinear dependencies among a set of variables once the nonlinear kernels have been properly defined in advance [53]. The feasibility of the kernel-based techniques relies on mapping the original image space to a higher-dimensional space without explicit knowledge of the functional mapping, which is typically known as the “kernel trick”. There are, however, two main limitations of the kernel-based techniques that may preclude a wider applicability in brain imaging. First, the accuracy and validity of the detected nonlinear dependencies are heavily linked to the chosen kernel, which typically depends on a previous knowledge of the datasets, limiting the search for more general and unknown associations. Second, since the functional mapping underlying the kernel trick is mostly unknown, it is not feasible to reconstruct the spatial loadings of the detected cross-correlated networks in the original space of the corresponding images (i.e., back projection to the original space). To overcome these limitations, several metrics like Distance Correlations/Covariances (DCC) [38,40] and the Hilbert-Schmidt independence criterion (HSIC) [55] have been proposed in the multivariate statistical analysis literature. As recently stated [41,56], HISC and DCC are strongly related and share similar formulations and many common properties. While the sample DCC is computed via pairwise Euclidean matrices defined for each dataset and the HSIC is computed via a Gaussian kernel, both metrics have been proven to be equal to zero if and only if the datasets are statistically independent [41]. In addition, even though distance correlation is equivalent to distance covariance for testing purposes, the former one has the advantage of being bounded by 1, as in the case of Pearson’s correlations. Further, the distance correlation is equal to one if and only if there is a linear relationship between the variables.

In this work, we take advantage of a recently proven equivalence between DCC and HSIC formulations. Such equivalence is based on a bijective transformation between Euclidean distances and kernels [41]. Hence, one can construct a distance-based kernel for testing independency between the datasets and use it in a more general context of Kernel CCA. In other words, we used an induced distance-based kernel to formulate a Canonical Distance Correlation Analysis (CDCA). Our choice is also justified by results in [55], showing that the maximum kernel canonical correlation for any generic kernel is also a valid measure of statistical independence.

From a computational perspective, our proposed CDCA is easy to implement. For each PET modality, a matrix of Euclidean distances between each pair of images (stacked as a high-dimensional vector) is first computed. Once the inter-distances matrices (denoted by *D*) have been computed, the corresponding kernels are constructed by following [41] as *K = 1-D/max(D)*. The eigenscores are then obtained by applying a classical KCCA with the previous distance-induced kernels. Notice that KCCA typically requires a regularization procedure [39,52,53] to maintain numerical stability and accuracy. In the current implementation, we selected the regularization parameter by employing a 10-fold cross-validation approach. The output from the CDCA is simply a set of score pairs (i.e., canonical distance-correlations) ordered according to their respective percent of explained co-variability. Thus, by definition, the CDC β-amyloid scores can be considered a spatial summary representation of the β-amyloid distribution that is maximally distance-correlated to certain (distributed) spatial network of tau.

### Voxelwise maps of distance-correlations

As in the case of the KCCA, the CDCA only allows us to extract pairs of scores that are maximally distance-correlated. By following kernel-based techniques, one would be able to reconstruct the so-called eigenimages of spatial loadings associated with each modality, but it is an extremely tedious and sometimes inaccurate process. Fortunately, the CDCA is based on a particular choice of the underlying kernel that is aimed at adapting to the concept of distance-correlation (i.e., distance-based induced kernel). Thus, the spatial loadings can be computed by performing a voxelwise distance-correlation analysis between each voxel in the image dataset and the corresponding CDCA scores. Notice that this procedure is similar to the typical intra-modality linear projection step used in SVD and CCA [51]. Hence, rather than using a linear projection via scalar products or Pearson’s correlations (e.g., images times scores operations), we associate images and scores via the distance-correlation metric.

As a result of the previous procedure, voxels with high spatial loading values co-vary together (i.e., are positively distance-correlated), while voxels with high opposite signed values are negatively distance-correlated. Thus, high spatial loadings of an eigenimage can be interpreted as a spatial network of highly distance-correlated voxels. Taken together, the ordered pairs of eigenimages produce the spatial representation of the CDCA scores that are maximally cross-distance-correlated. In an analogous manner, the CDCA scores in one modality can be distance-correlated to each of the voxels in the other modality (i.e., similar to a voxelwise linear regression analysis) to produce distributed-to-distributed views of full cross-distance-correlation between the two modalities. For simplicity, the images resulting from such cross-distance-correlation analysis will be referred to as cross-eigenimages maps. Additionally, one can compute a so-called homologous (i.e., local) distance-correlation image via the distance-correlation metric between corresponding voxels in the two image modalities. Note that such homologous distance-correlation would produce a local-to-local (at the voxel level) distributed view of the (nonlinear) associations between β-amyloid and tau PET SUVR images.

In this study, the voxelwise maps of cross-distance-correlations were thresholded by employing permutation techniques to derive the empirical distribution of the maximum (over the space of voxels) distance-correlation, which is a common approach to control for multiple comparisons [57].

## Results

The subject characteristics analysis from Table 1 revealed statistically significant associations between the clinical classification and APOE ε4 status (*χ^2^*=5.89, *p*=0.05) and gender (*χ^2^*=9.85, *p* = 0.007), but not age (*F* = 1.24, *p*=0.29). The MMSE (*F* = 2.01, *p*=0.11) measure showed no statistically significant differences based on the clinical classification, but the ADAS-Cog (*F* = 9.36, *p*<0.001) did.

**Table 1.** Summary of subject characteristics.

|  | **All** | **CN** | **EMCI** | **MCI** |
| --- | --- | --- | --- | --- |
| **Sample Size** | 157 | 28 | 13 | 116 |
| **Age** | 71.90 ± 7.79 | 69.66 ± 6.08 | 70.60 ± 8.58 | 72.61 ± 8.01 |
| **Gender (F/M)** | 78/79 | 20/8 | 9/4 | 49/67 |
| **APOE ε4 (Carrier/Non-Carrier)** | 33/124 | 10/18 | 4/9 | 19/97 |
| **Aβ (High / Low)** | 90/67 | 17/11 | 5/8 | 68/48 |
| **MMSE** | 28.17 ± 2.23 | 28.84 ± 2.15 | 29.08 ± 1.31 | 27.91 ± 2.29 |
| **ADAS-Cog** | 11.68 ± 4.97 | 9.13 ± 6.11 | 6.80 ± 3.17 | 12.82 ± 4.29 |

The first, second, and third canonical distance-correlations components accounted for 26.31%, 8.61%, and 6.30% of the total co-variability between the florbetapir and flortaucipir PET images, respectively. Figures 1A and 1B show the spatial representation of the corresponding CDC loadings for each PET modality. The strongest positive weights in the first β-amyloid eigenimage (Figure 1A) correspond to the medial prefrontal and posterior cingulate cortices, precuneus, lateral inferior temporal, and fusiform gyri, although they appear spread over the cortex. Correspondingly, the strongest positive weights in the first tau eigenimage (Figure 1B) cover areas of the lateral inferior temporal gyri and the bilateral fusiform gyrus, following a remarkable resemblance to the proposed topography of Braak’s stage III/IV of tau accumulation [3]. It is worth noting that the CDC eigenimages do not necessarily represent spatial networks with high accumulations of either β-amyloid or tau. They are simply spatial loadings contributing to the corresponding scores that are maximally distance-correlation according to the CDCA. Figure 1C shows the scatter plot for the β-amyloid and tau first CDC scores, where each color represents a different cognitive subpopulation. That pair of scores produced a maximum distance-correlation value of *dc*=0.49 (*t*=54.27, *p*<0.001), which is evidence that the β-amyloid and tau scores are not statistically independent values, but follow a general (possible nonlinear) dependency. The first β-amyloid CDC scores were highly correlated to Statistical ROI (Stat-ROI) measurements (*r=*0.97, *dc*=0.95). Indeed, it is a clear indication that the CDCA-based β-amyloid scores do not only account for a significant relationship with distributed tau uptake, but can also be considered a feasible biomarker to segregate between Aβ_L_ and Aβ_H_ subjects. Figure 1D shows boxplots corresponding to the individual β-amyloid and tau CDC scores for the three different clinical classification groups. The mean values for the β-amyloid CDC scores were SUVR = 1.27 ± 0.29, 1.20 ± 0.25, and 1.35 ± 0.34 for the CN, EMCI, and MCI groups, respectively. These values resulted in no statistically significant differences across the clinical classification (*F* = 1.22, *p*=0.30) and very weak distance-correlations were found between the CDCA β-amyloid scores and MMSE (*dc*=0.02) and ADAS-Cog (*dc*=0.06). On the other hand, the mean values for the tau CDCA scores were SUVR = 1.05 ± 0.05, 1.06 ± 0.05, and 1.09 ± 0.09, and were not statistically significant different (*F* = 2.34, *p*=0.07) across the CN, EMCI, and MCI groups. The distance-correlations between the CDC tau scores and the MMSE and ADAS-Cog scores were *dc*=0.08 and *dc*=0.15, respectively, which are still weak values for stablishing some sort of dependency, but they are certainly higher than the ones obtained for the CDC β-amyloid scores.

**Figure 1.**
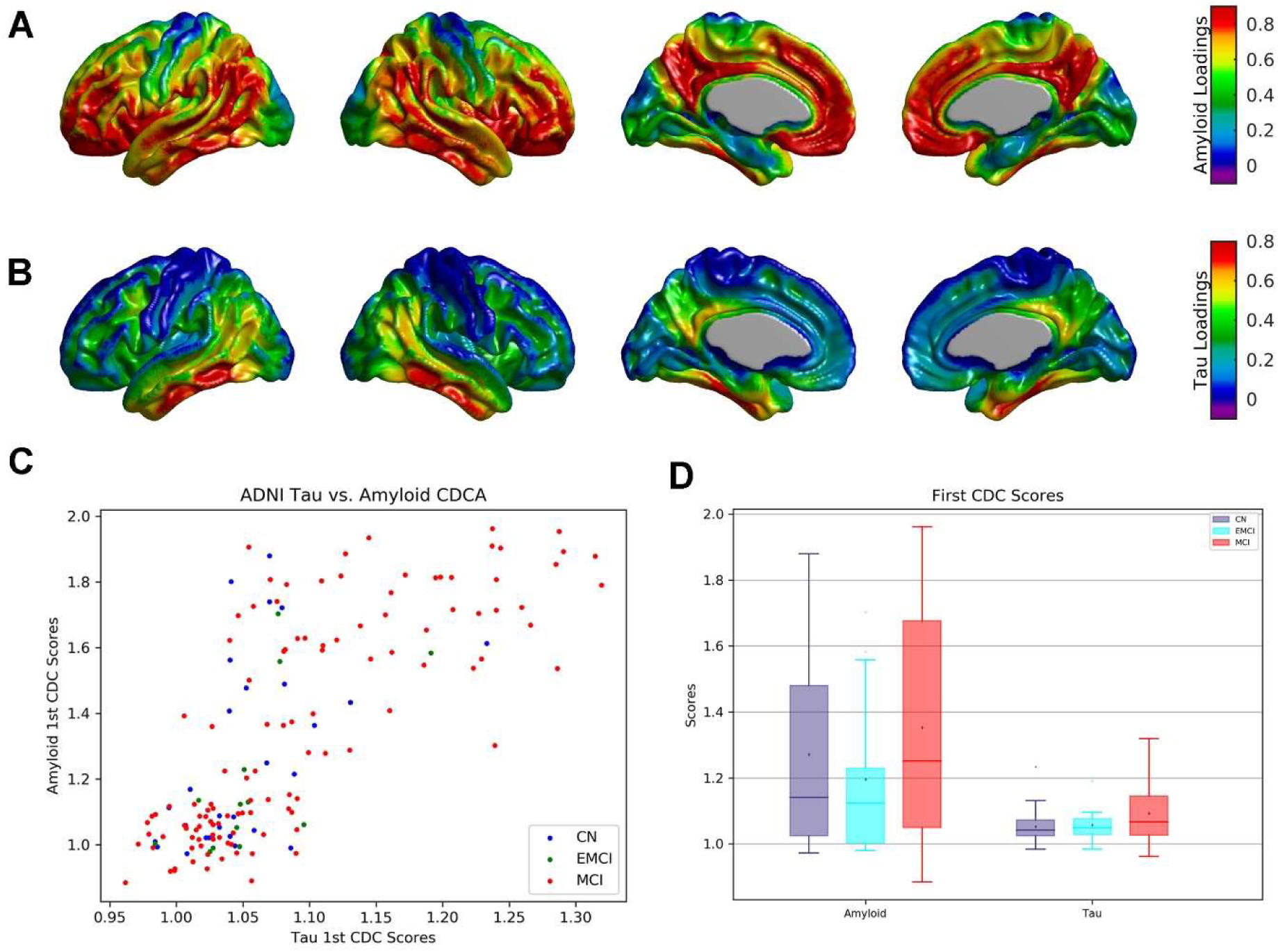
CDCA between β-amyloid and tau across the whole sample of florbetapir and flortaucipir PET images. (A, B) Cortical surface projections for the pair of spatial loadings of (A) β-amyloid and (B) tau corresponding to the first canonical correlation. The strongest positive weights in (A) and (B) are regions maximally distance-correlated with CDC β-amyloid and tau scores, respectively. (C) Scatter plot between the resulting β-amyloid and tau scores across each subject’s cognitive classification. (D) Boxplots of the β-amyloid and tau scores for each cognitive group. No statistically significant differences were observed across clinical classification.

Figure 2 presents the thresholded distributed-to-distributed views of the β-amyloid-distributed tau and tau-distributed β-amyloid associations, respectively. Here, the distributed nature of the β-amyloid and tau measurements comes from the fact that they have been previously derived as CDC scores that are simply a spatial summary representation of the β-amyloid and tau uptake in distributed networks that are maximally distance-correlated. Since this cross-distance-correlation analysis is similar to a voxelwise regression, the linear effect of age, sex, and the ADAS-Cog score have been removed from both datasets and the CDC scores before computation. The strongest, statistically significant, positive weights (threshold = 0.131) in the first β-amyloid cross-distance-correlations map (Figure 2A) correspond to the medial prefrontal and posterior cingulate cortices, precuneus, and lateral inferior temporal regions. In other words, those are the spatially distributed β-amyloid regions that are maximally distance-correlated with the (distributed) tau scores. Correspondingly, the widely spread distribution of β-amyloid represented by the β-amyloid scores is maximally distance-correlated (threshold = 0.154) to tau in the entorhinal cortex (Figure 2B), a clear resemblance with the topography of Braak’s stage I/II.

**Figure 2.**
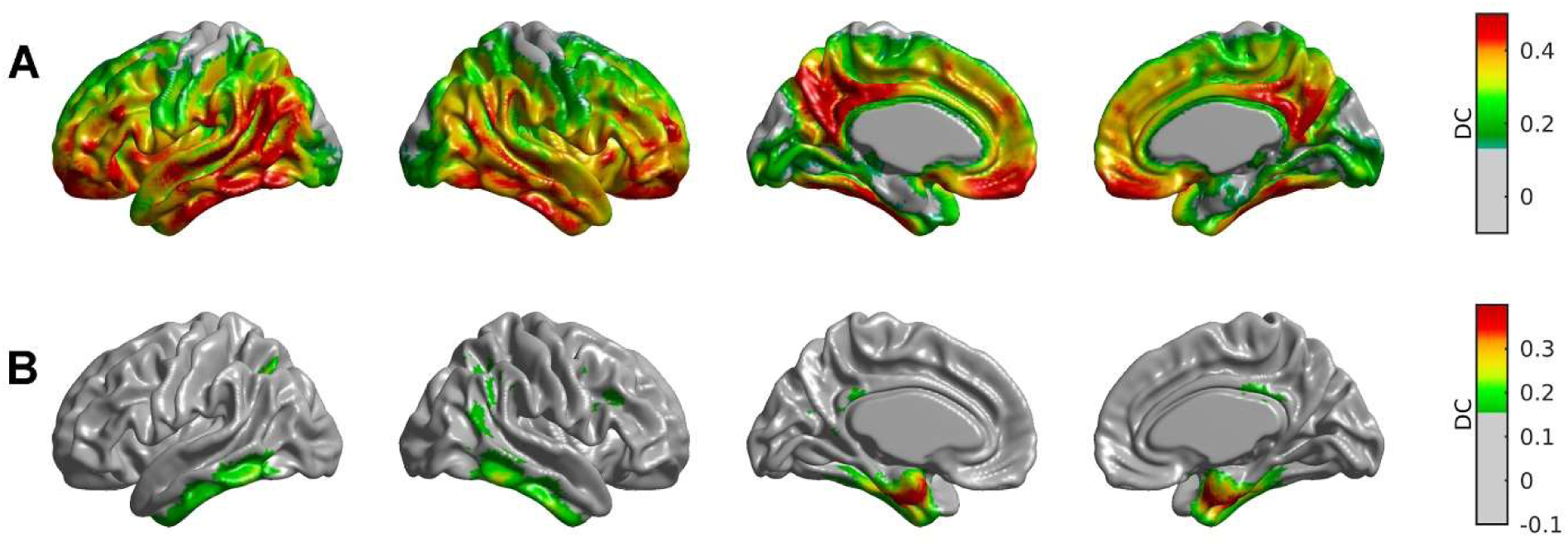
Statistical assessment of distance-correlations cross-eigenimages of amyloid and tau. Cortical surface projections of thresholded maps of distance-correlations using permutation techniques. The β-amyloid cross-eigenimage (A) appeared to be statistically significant for large areas of the cortex, particularly in the temporal lobe, posterior cingulate, and medial frontal cortex. The statistically significant regions for the tau cross-eigenimage (B) appeared more localized to the entorhinal cortex and the lateral inferior temporal gyri.

The CDCA produced just a single distributed-to-distributed view of the β-amyloid-tau relationship among the possible multiple views that could be derived from such an association. In fact, the most common distributed-to-distributed view could be obtained from the so-called global-to-distributed cross-correlated view that is derived from the global average of the β-amyloid and tau SUVR measurements. Hence, by computing distance-correlations of these global measurements, we obtained the global tau-distributed β-amyloid and global β-amyloid-distributed tau spatial representations shown in Supplementary Figure 1. There, the global β-amyloid measurements (Supp. Figure 1B) produced a similar distance-correlations pattern with tau (i.e., localized in the entorhinal cortex) as the one obtained with the CDC β-amyloid scores (see Figure 2B). However, the global tau measurements produced very weak distance-correlations with β-amyloid (see Supp. Figure 1A) as compared with the CDC tau scores (see Figure 1A), reinforcing the idea that the β-amyloid-tau associations follow a very distinctive localized tau – globally-spread β-amyloid pattern.

The CDCA was also re-assessed after the inclusion (in separate models) of APOE ε4 genotype, entorhinal cortical thickness, and hippocampal volume as potential confounding variables that may be mediating the spatial association between PET β-amyloid and tau. The resulting cross-eigenimages maps from this analysis are presented in Supplementary Figure 2. Notice that the removal of APOE ε4 genotype seems to slightly decrease the intensity DC values of the tau cross-eigenimage, as compared to Figure 2B (no removal of APOE ε4). However, the removal of the effect of either cortical thickness in the entorhinal cortex or hippocampal volume preserves the strength and distribution of the DC values for the tau cross-eigenimage. In contrast, the removal of the effect of these three confounding variables seems to decrease the DC values for the β-amyloid cross-eigenimage, particularly within the lateral temporal-parietal regions and the medial frontal cortex.

We subsequently re-evaluated the CDCA for the subpopulations of CN and MCI subjects. The group of EMCI was not reassessed due to its relatively low sample size. First, a voxelwise regression analysis was performed by including β-amyloid or tau uptake as dependent variables and cognitive group (CN and MCI) as a regressor of interest. Such analysis revealed that there were no statistically significant differences between MCI and CN for either β-amyloid or tau uptake. The resulting t-SPMs (Statistical Parametric Maps) for the contrast MCI-CN are shown in Supplementary Figure 3. In both cases, no voxel in the cortex survived multiple comparisons thresholding (using False Discovery Rate) at 0.05 level. The CDCA revealed that the first canonical component accounted for 29.41% and 28.90% of the total co-variability in the CN and MCI groups, respectively. The first pair of canonical components produced distance-correlation values of *dc*=0.81 (*t*=16.45, *p*<0.01) and *dc*=0.55 (*t*=44.47, *p*<0.01) for the CN and MCI groups, respectively. As in the case of the whole sample, the first β-amyloid CDC scores were highly correlated to Stat-ROI measurements in both the CN group (*r=*0.96, *dc*=0.86) and the MCI group (*r=*0.98, *dc*=0.95). Figure 3 shows the first pair of canonical cross-distance-correlation maps for β-amyloid and tau. Notice that the β-amyloid cross-eigenimage seems to be strikingly stronger in CN than in MCI, where the loadings cover extended areas of the medial prefrontal cortex, posterior cingulate cortex, and precuneus. In contrast, the tau cross-eigenimage carry stronger loads in MCI than in CN, particularly within the entorhinal cortex. To eliminate a potential bias due to the relatively low sample size in the CN group (n=28) as compared to the MCI group (n=116), we re-evaluated the CDCA by means of a random sampling analysis in MCI group. We performed 100 random shufflings of the n=116 samples in the MCI group and for each shuffling, we selected a random subsample of n=28 MCI subjects that were submitted to a CDCA. Finally, we computed the average of the spatial β-amyloid and tau cross-eigenimages across the 100 repetitions, which are presented in Supplementary Figure 4. Notice that the tau cross-eigenimage values within the entorhinal cortex remain almost identical to the ones corresponding to the whole MCI sample (see Figure 3D), although some strong distance-correlations values also appear in the lateral inferior temporal gyri and fusiform gyri. On the other hand, the β-amyloid cross-eigenimage values appear to be stronger during the random sampling procedure, particularly in the precuneus and the posterior cingulate cortex. To summarize, the random sub-sampling analysis produced β-amyloid and tau cross-eigenimage maps with stronger values as compared to the whole MCI sample. The difference in cross-correlation spatial patterns between CN and MCI groups is evident even with the sub-sampling procedure, which is a compelling argument to claim that the spatial dependency between β-amyloid and tau seems to follow distinctive patterns that depend on the cognitive status.

**Figure 3.**
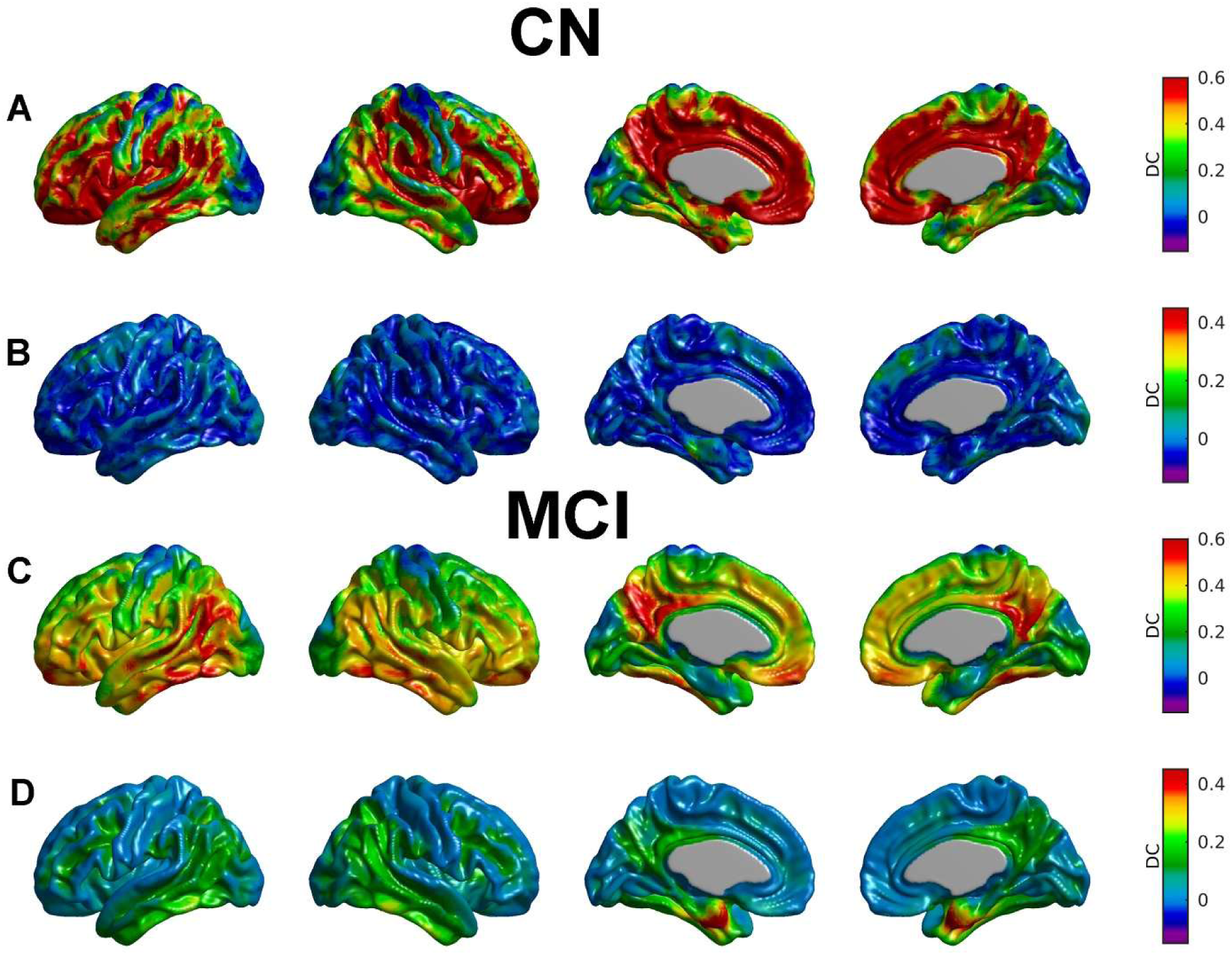
CDCA between β-amyloid and tau across each subsample of CN and MCI subjects. The distance-correlation values in the β-amyloid cross-eigenimages (A and C) appeared to be stronger and more spatially spread for the CN sample (A) than for the MCI subjects (C). Strong values for the tau cross-eigenimage in the entorhinal cortex appeared only relevant to the MCI subjects subsample (B and D).

The CDCA was also assessed for the aggregation of subjects according to their β-amyloid status according to their uptake in the Stat-ROI (i.e., low and high β-amyloid). The first canonical component accounted for 11.75% and 16.75% of the total co-variability in the Aβ_L_ and Aβ_H_ groups, respectively. The first β-amyloid CDC scores were weakly distance-correlated to the Stat-ROI SUVR values in the Aβ_L_ group (*r=*0.59, *dc*=0.36), but highly distance-correlated for the Aβ_H_ group (*r=*0.95, *dc*=0.88). Figure 4 shows the first β-amyloid and tau cross-eigenimages for the Aβ_H_ and Aβ_L_ groups. The β-amyloid cross-eigenimage shows stronger distance-correlations values for the Aβ_H_ group, particularly in areas covering the lateral temporal gyri (Figure 4C). In contrast, the tau cross-eigenimage corresponding to the Aβ_L_ group produced low distance-correlation values, which is an indication that the process of β-amyloid accumulation with minimal burden seems to be statistically independent of the distributed tau uptake.

**Figure 4.**
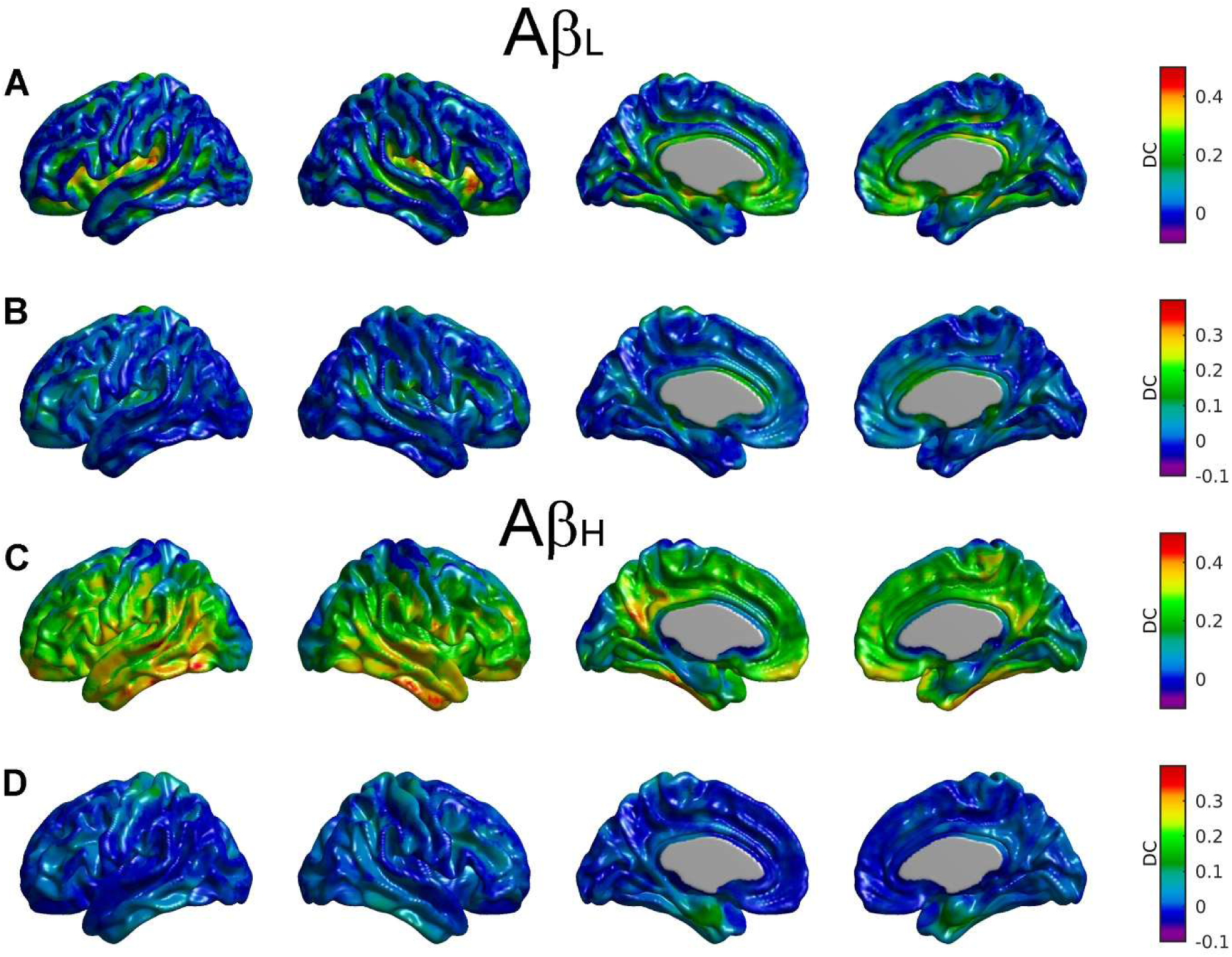
Statistical assessment of CDCA for the cohort of Aβ_L_ and Aβ_H_ subjects. The distance-correlation values in the β-amyloid cross-eigenimages (A and C) are stronger and more spatially spread for the Aβ_H_ subjects (C). In contrast, the DC values in the entorhinal for the tau cross-eigenimages (B and D) are very similar, where only small, subtle increases are observed for the Aβ_H_ cohort.

However, the tau loadings for the Aβ_H_ groups produced relatively stronger values in the entorhinal cortex, although not as elevated as in the case of the whole sample. This finding is an indication that the spatial relationship between β-amyloid and tau is not totally determined by the amount of extreme (i.e., very low or very high) β-amyloid uptake, but follows a continuously evolving dependency that reaches its maximal expression at intermediate stages of β-amyloid accumulation.

Finally, Figure 5 shows the maps of the local-to-local distance-correlations between β-amyloid and tau for the whole sample as well as for the sub-samples of CN and MCI subjects. The strongest local distance-correlations for the whole sample analysis appeared in the bilateral precentral and postcentral gyrus, superior parietal lobule, and to a lesser extent in the lateral inferior temporal gyri, fusiform gyri, and small portions of the entorhinal cortex (Figure 5C). However, the local distance-correlations in the lateral inferior temporal gyri and entorhinal cortex only appear to be significantly strong for the group of MCI subjects (Figure 5B).

**Figure 5.**
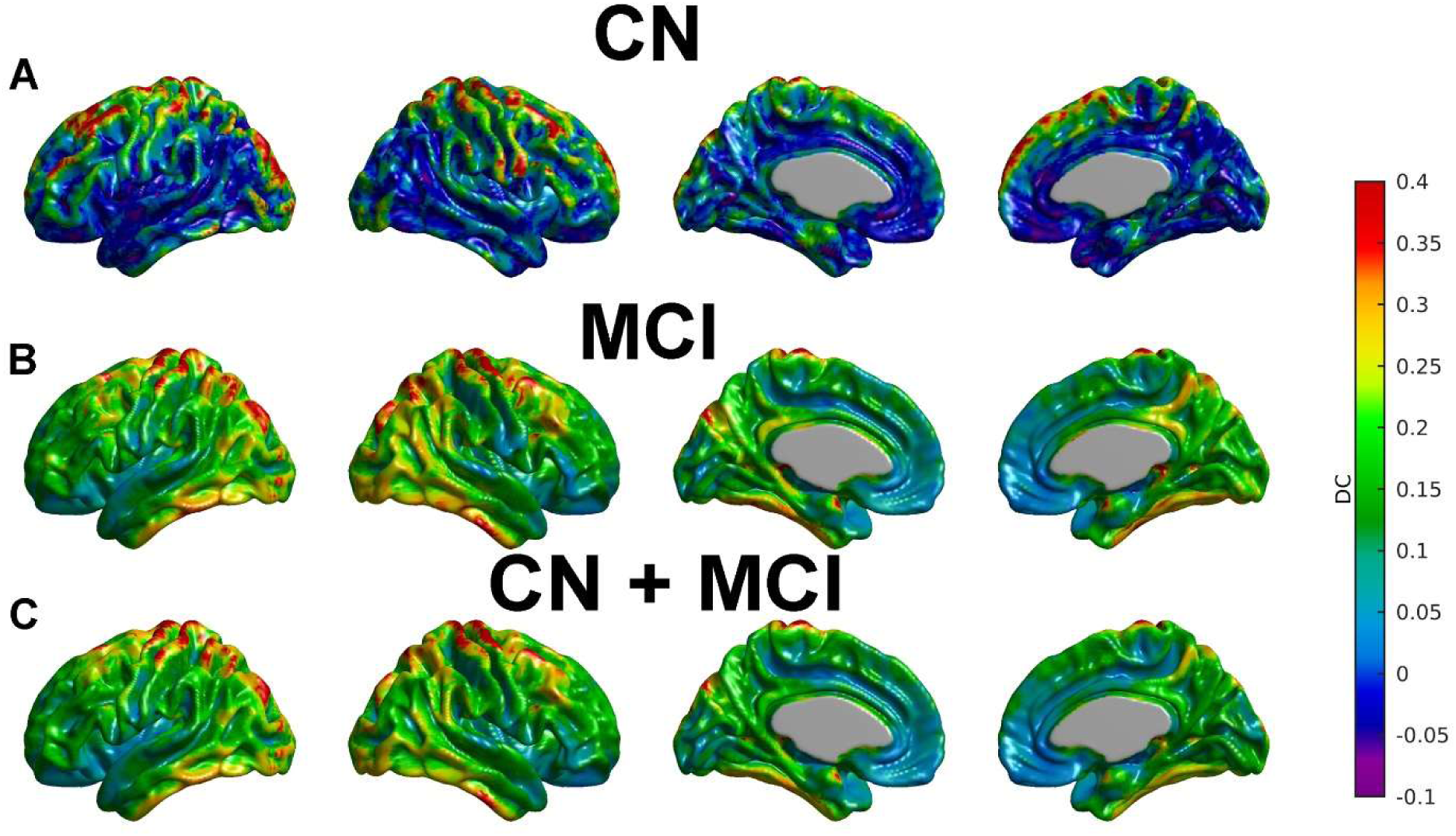
Local distance-correlation analysis for different subsamples of subjects. The overall dependency between local measurements of β-amyloid and tau seems to be stronger for MCI subjects (B). Such dependency in the whole sample seems to be driven by the MCI subsample (C).

## Discussion

We have proposed a Canonical Distance Correlation Analysis that allowed us to reveal not only multi-modality linear interactions, but also more complex nonlinear associations between β-amyloid and tau PET data. The CDCA approach introduced here generalizes our previous SVD cross-correlation analysis [51] in the sense that it can detect linear as well as non-linear cross-modalities dependencies. Since our approach is ultimately based on a kernel CCA, it is possible to fully characterize the full cross-correlation matrix at the imaging voxel level with just a few pairs of canonical components that contain most of the explained inter-modalities co-variability. Our approach provides a unified framework to uncover distributed-to-distributed, local-to-distributed, global-to-distributed, and local-to-local views of inter-modality associations that might be related by underlying complex and possible nonlinear dependencies.

A first attempt to construct a kernel-based canonical correlation analysis relying on distance-correlations was proposed in [58]. Even though the authors in [58] constructed a kernel using the distance-correlations metric, their method cannot be regarded as a kernel CCA approach. Indeed, their method operates in the space of the image features rather than in the space of the samples, which is the common choice in kernel-based approaches and in the CDCA introduced here. This approach is a limitation from a computational and practical point of view since the image features space might comprise the totality of the image voxels as is typical in voxelwise CCA. In contrast, the CDCA introduced here employs an induced distance-based kernel defined in the samples space, with is typically of much lower dimension than the image features space.

Additionally, our method used a natural approach based on distance-correlations to recover the spatial representation of the canonical distance-correlations scores.

It has been a common practice to assess the spatial relationship between β-amyloid and tau PET images at the local level [35–37]. Such practice may originate in early efforts for proving that β-amyloid and tau act in a coordinated, synergistic manner that leads to neurodegeneration and cognitive decline. Thus, [37] and [36] found significant local correlations between β-amyloid and tau in the lateral inferior temporal lobe for respective samples of cognitively normal older subjects. Using a small sample of clinically probable AD subjects, [35] found strong local-to-local correlations in small regions of the paracentral lobule, lateral middle frontal, and superior parietal cortices. Our findings about local-to-local associations are consistent with those of [35–37]. However, we found that the local-to-local distance-correlations in the lateral inferior temporal gyri were significantly stronger in the case of MCI subjects as compared to the CN group. On the other hand, the local-to-local distance correlations corresponding to the paracentral lobule and superior parietal cortex remain consistent across the two different cognitive groups.

The local-to-local correlational view has been consistently complemented with more general approaches to cover the so-called local-to-distributed view and, to some extent, the distributed-to-distributed (network) view of the β-amyloid-tau relationship. While the local-to-distributed view is relatively easy to implement by means of the seed-based correlation approach, the distributed-to-distributed view requires a more complex analysis of full cross-correlation matrices that may result in computationally prohibitive burden at the voxelwise scale. Hence, [36] computed a full cross-correlation matrix between β-amyloid and tau PET measurements of CN older adults at the ROI spatial scale to conclude that there were strong associations between β-amyloid uptake across multiple regions in the brain and tau uptake in the bilateral temporal neocortex. Using the same ROI cross-correlation matrix approach in a small cohort of subjects with probable AD dementia, [35] found weak (i.e., not statistically significant after multiple comparisons correction) associations between β-amyloid uptake in the parietal and occipital cortices and tau accumulation in the lingual gyri, cuneus, and occipital areas. Also using an ROI approach, [30] performed a canonical correlation analysis to uncover the distributed-to-distributed view in the β-amyloid-tau relationship in a small sample of CN and mild AD subjects. It was found that the tau topography was predominantly strong in the medial temporal lobe, parietal cortex, and precuneus, and it was maximally correlated to the β-amyloid topography with high loads in the frontal and parietal regions [30]. In a more general scenario of correlation analysis at the imaging voxel scale, [37] used a weighted degree of connectivity measure to summarize the full cross-correlation voxel-by-voxel matrix between imaging-derived β-amyloid and tau measures in CN subjects, which also allowed an automatic way of defining inter-connectivity hubs for both modalities. Thus, [37] found that areas including the inferior-lateral temporal cortex and entorhinal cortex yielded a significant elevated number of tau correlations with distributed β-amyloid. Correspondingly, areas such as the lateral-ventral frontal, inferior-parietal, and lateral temporal cortices showed many significant associations between β-amyloid and distributed tau [37]. Our kernel-based CDCA revealed a distinctive spatial association pattern between β-amyloid and tau that appears to vary according to the cognitive stage. The first β-amyloid canonical DC that produced a maximal distance-correlation with distributed tau turned out to be highly correlated with a Stat-ROI measurement that discriminates between low and high levels of β-amyloid uptake. This finding was valid not only for the whole sample of CN and MCI subjects, but it also held for the two individual cognitive sub-samples as well. This result is in correspondence with recent findings of [59] that showed that 18F-flortaucipir PET was able to discriminate amyloid-positive from amyloid-negative forms of neurodegeneration. Thus, in principle, one might interpret that the distributed β-amyloid CDC scores were just a reflection of the slow progression from low to high levels of β-amyloid accumulation. However, Stat-ROI SUVR measurements and the β-amyloid scores were highly distance-correlated for the Aβ_H_ group only. This finding is an indication that the CDCA may be generating scores that account for a more complex underlying dependency between β-amyloid and tau than a mere reflection of the progression of β-amyloid accumulation. Additionally, in correspondence with [30] and [37], our CDC distributed tau scores produced high values of distance-correlations with β-amyloid that were spread over extended areas of the cerebral cortex. It was interesting to note that such an association pattern was achieved with relatively low loads of distributed tau in the case of CN subjects. In contrast, the tau loads were particularly strong in the inferior temporal regions in the MCI subjects. On the other hand, the β-amyloid loads cover typical regions of known AD-related vulnerability for both CN and MCI groups, but produce a distinctive pattern of distance-correlations with tau. Indeed, strong correlations between β-amyloid scores and tau were observed only for the MCI group and were highly localized in the entorhinal cortex. Interestingly, this dependency pattern does not seem to be determined by a high amyloid uptake condition since the spatial relationship between distributed β-amyloid and tau in the entorhinal cortex was not necessarily strong for the Aβ_H_ group.

Taken together, our results suggest that the distributed β-amyloid – localized tau dependency is not completely determined by the β-amyloid accumulation process at the initial stages of AD. A possibility is that the spatial patterns of strong cross-correlations between widespread β-amyloid and localized tau in the entorhinal cortex and lateral inferior temporal lobe might be mediated by underlying molecular mechanisms and/or genetic factors with a more direct influence on the cognitive deterioration during the AD progression. Our analysis showed (see Supplementary Figure 2) that the inclusion of APOE ε4 genotype does not substantially modify such cross-correlation patterns. Similarly, well-known biomarkers of neurodegeneration, such as entorhinal cortical thickness and hippocampal volume, do not seem to confound the association between tau and β-amyloid PET for the whole sample of CN and MCI. However, such analysis was only confined to global metrics of neurodegeneration with a very limited spatial specification. A more exhaustive analysis involving other neurodegeneration biomarkers, such as FDG PET, could produce a more detailed and comprehensive explanation about possible factors mediating the β-amyloid-tau relationship, but it is beyond the scope of the current study.

## Conclusions

We uncovered spatial associations between tau and β-amyloid in a sample of CN and MCI subjects. By exploring the large-scale cross-distance-correlation between β-amyloid and tau PET images with the CDCA approach, our analysis revealed a very distinctive pattern of distributed β-amyloid – localized tau that seems to vary according to cognitive status. More importantly, such spatial dependency does not rely on low or high levels of β-amyloid accumulation or the differences in β-amyloid and tau uptakes between cognitively normal and mild cognitive impairment conditions. To our knowledge, this study is the first to relate tau and β-amyloid burden from a nonlinear network perspective that accounts for distributed-to-distributed and local-to-distributed patterns of cross-distance-correlation. The distinctive cognition-varying amyloid-tau relationship revealed by our CDCA could have important implications for the population enrichment of clinical trials. It has been shown that the specific co-variation patterns in the amyloid-tau relationship can distinguish between CN and MCI despite the fact that their individual uptakes are unable to do so. Thus, using the CDC scores might be more accurate than the amyloid or tau SUVR for the enrollment in clinical trials of those individuals in the path of cognitive deterioration. Future work will also expand the current multivariate analysis to identify either distributed or local patterns of the β-amyloid-tau association that are maximally related (e.g., modulated) to hypometabolism and/or metabolic connectivity.

## Acknowledgments

Data collection and sharing for this project was funded by the Alzheimer’s Disease Neuroimaging Initiative (ADNI) (National Institutes of Health Grant U01 AG024904) and DOD ADNI (Department of Defense award number W81XWH-12-2-0012). ADNI is funded by the National Institute on Aging, the National Institute of Biomedical Imaging and Bioengineering, and through generous contributions from the following: AbbVie, Alzheimer’s Association; Alzheimer’s Drug Discovery Foundation; Araclon Biotech; BioClinica, Inc.; Biogen; Bristol-Myers Squibb Company; CereSpir, Inc.; Cogstate; Eisai Inc.; Elan Pharmaceuticals, Inc.; Eli Lilly and Company; EuroImmun; F. Hoffmann-La Roche Ltd and its affiliated company Genentech, Inc.; Fujirebio; GE Healthcare; IXICO Ltd.; Janssen Alzheimer Immunotherapy Research & Development, LLC.; Johnson & Johnson Pharmaceutical Research & Development LLC.; Lumosity; Lundbeck; Merck & Co., Inc.; Meso Scale Diagnostics, LLC.; NeuroRx Research; Neurotrack Technologies; Novartis Pharmaceuticals Corporation; Pfizer Inc.; Piramal Imaging; Servier; Takeda Pharmaceutical Company; and Transition Therapeutics. The Canadian Institutes of Health Research is providing funds to support ADNI clinical sites in Canada. Private sector contributions are facilitated by the Foundation for the National Institutes of Health (www.fnih.org). The grantee organization is the Northern California Institute for Research and Education, and the study is coordinated by the Alzheimer’s Therapeutic Research Institute at the University of Southern California. ADNI data are disseminated by the Laboratory for Neuro Imaging at the University of Southern California.

## Author contributions

FC and BJB designed research; FC, CM, APZ, and BJB performed research; FC and BJB analyzed the data and drafted the paper.

## Compliance with ethical standards

### Ethical approval

All procedures performed in studies involving human participants were in accordance with the ethical standards of the institutional and/or national research committee and with the 1964 Helsinki declaration and its later amendments or comparable ethical standards.

### Informed consent

Informed consent was obtained from all individual participants included in the study.

---

**Supplementary Figure 1.**
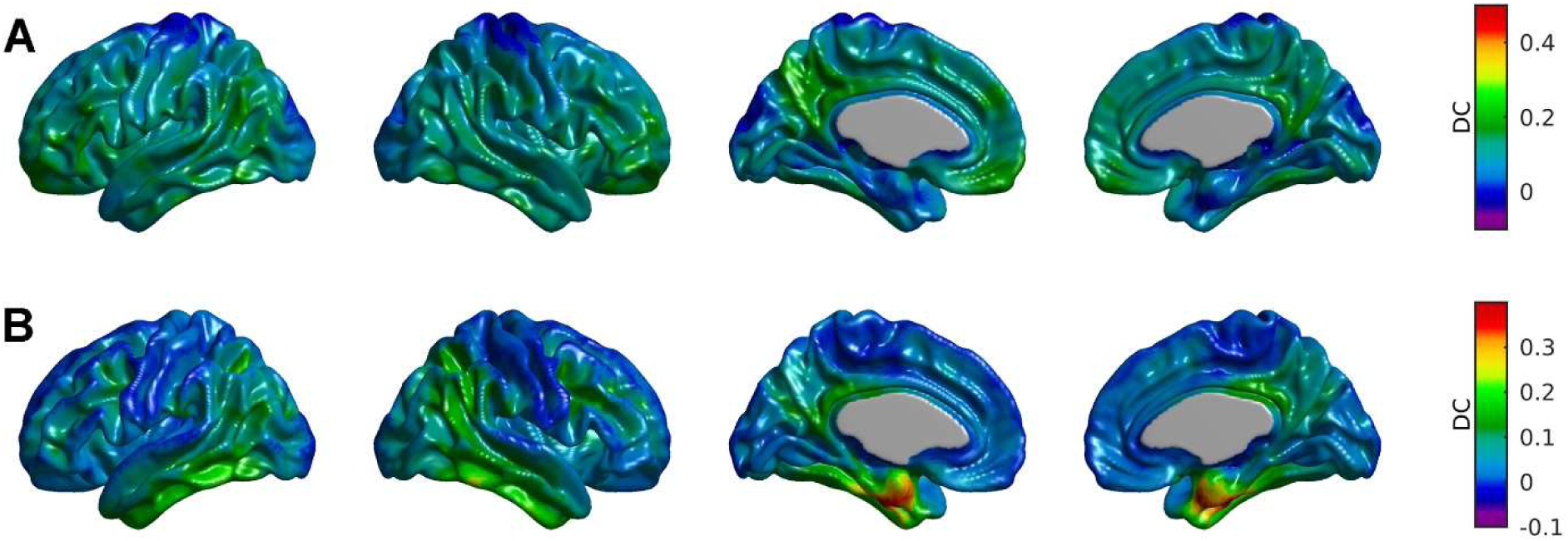
Statistical assessment of cross-eigenimages using the global measurements of β-amyloid and tau rather than the CDC scores. The β-amyloid cross-eigenimage produced by the global tau measurements (A) does not appear to strong and spatially spread as in the case of the tau CDC scores. In contrast, the tau cross-eigenimage produced high values of distance-correlations within the entorhinal cortex.

**Supplementary Figure 2.**
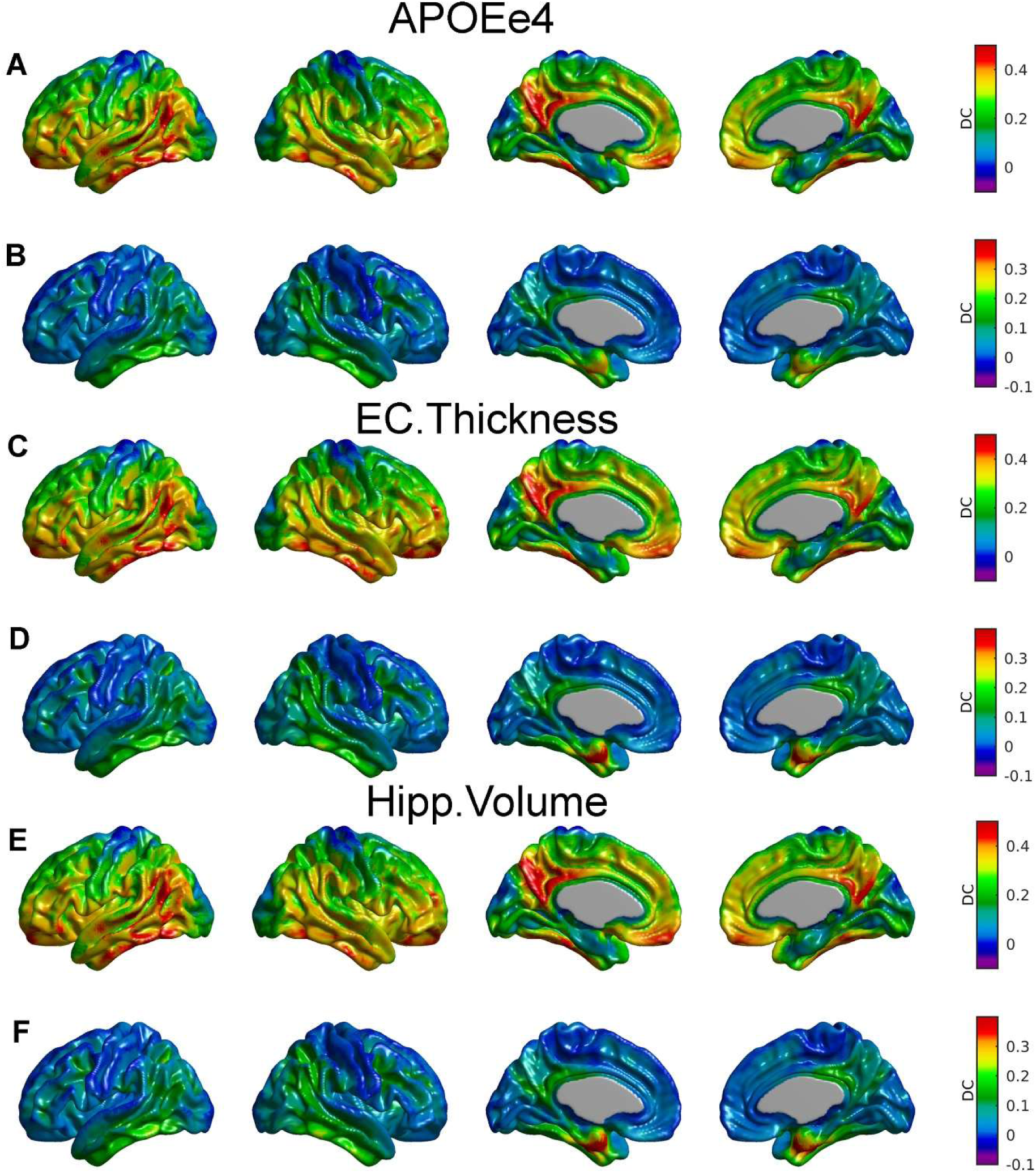
Statistical assessment of the CDCA between β-amyloid and tau PET images after the removal of the effect of ApoE ε4 genotype (A, B), entorhinal cortical thickness (C, D), and hippocampal volume (E, F). There was a small overall decrease in DC values for the β-amyloid cross-eigenimages (A, C, E) as compared to the original CDC analysis (Figure 2). The DC values for the tau cross-eigenimage in the entorhinal cortex appears to remain strong after the removal of the effect of the entorhinal cortical thickness and hippocampal volume (D, F). Only an evident decrease in DC within the entorhinal cortex for the tau cross-eigenimage was observed after the removal of the ApoE ε4 genotype effect (B).

**Supplementary Figure 3.**
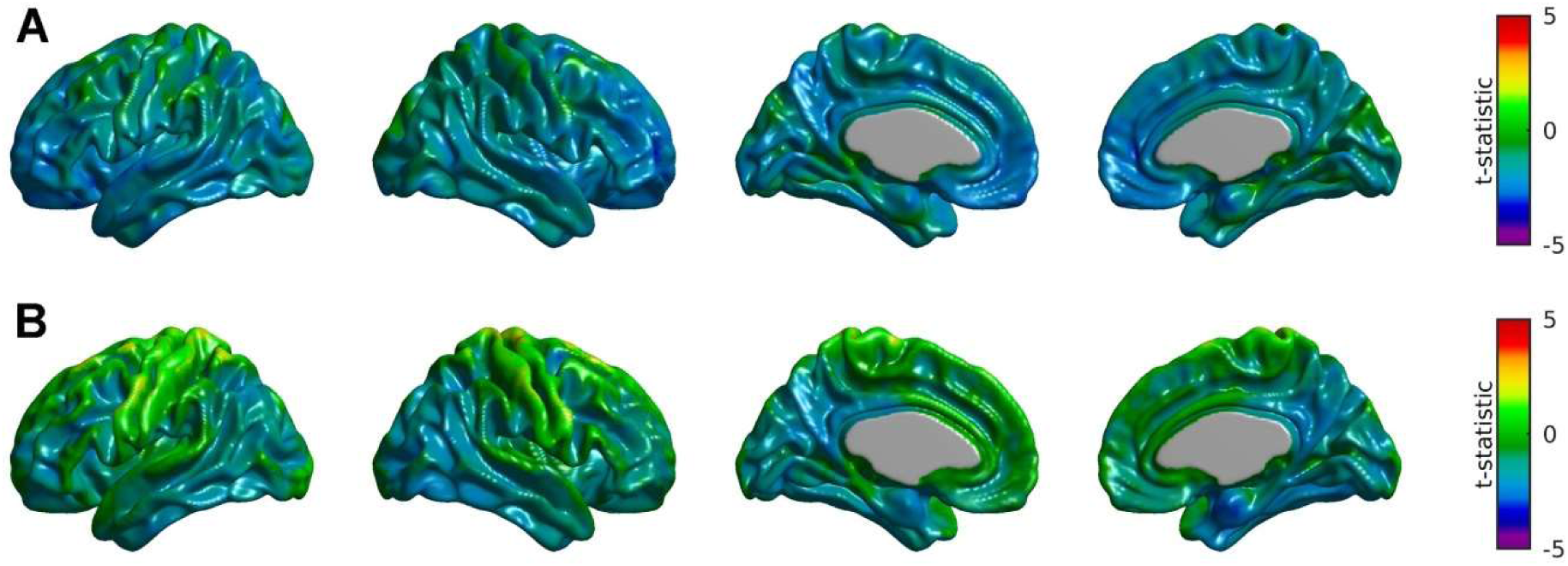
Statistical comparison of β-amyloid and tau uptakes between CN and MCI subjects. For both cases, FDR thresholding revealed no regions of statistically significant differences.

**Supplementary Figure 4.**
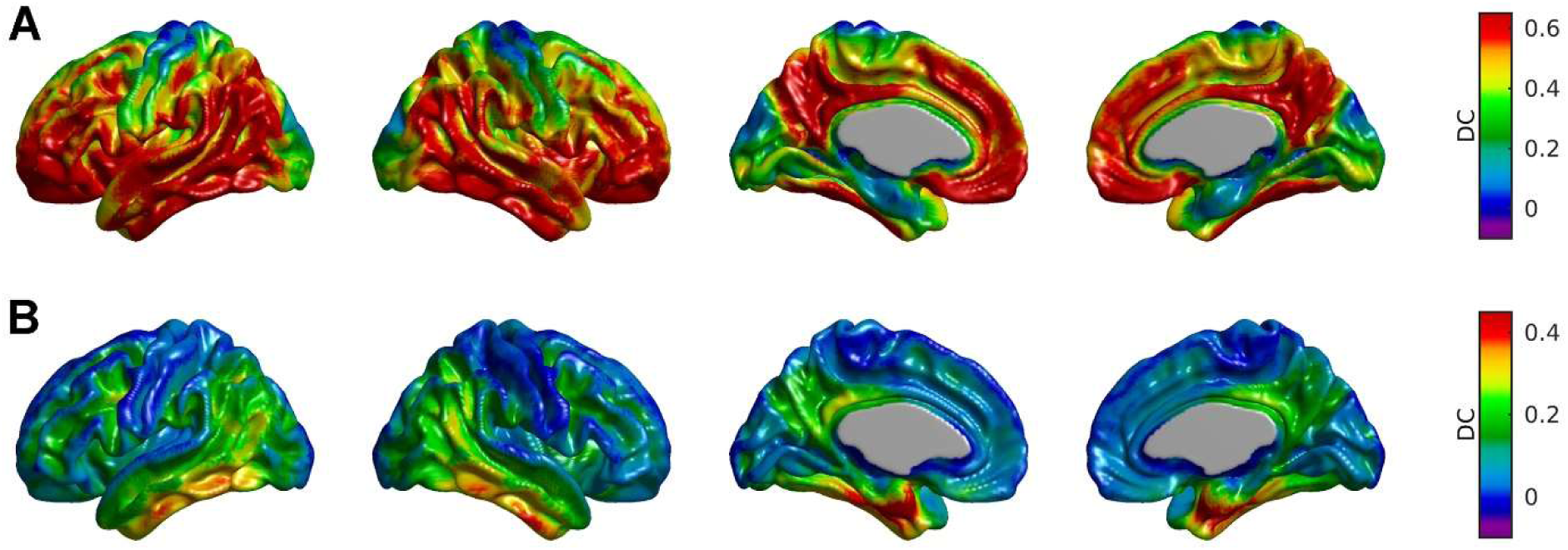
Statistical assessment of CDCA after 1000 repetitions with a subsample of N=28 MCI subjects. The main differences between CN and MCI spatial patterns of cross-eigenimages remain after the re-sampling analysis in the MCI subsample.

